# Neutrophil extracellular traps and thrombotic components across human coronary plaque progression: Association of intraplaque hemorrhage with NET formation

**DOI:** 10.1101/2024.06.10.598167

**Authors:** Eriko Nakamura, Saki Horiuchi, Kazunari Maekawa, Eiteru Kuroki, Toshihiro Gi, Nobuyuki Oguri, Murasaki Aman, Sayaka Moriguchi-Goto, Yujiro Asada, Atsushi Yamashita

## Abstract

Neutrophil extracellular traps (NETs), fibrin, and von Willebrand factor (VWF) are present in acute coronary thrombotic plaques. This study aimed to investigate their distribution among the coronary atherosclerotic progression and the potential association between intraplaque hemorrhage and NET formation.

Coronary artery sections from autopsy cases of non-cardiac death (n = 5, 37 sections) and ischemic heart disease (n = 11, 65 sections) were classified into six morphological groups and examined immunohistochemically. Effects of intraplaque hemorrhage related molecules on NETs formation were examined with human peripheral blood derived neutrophils.

Neutrophils and citrullinated histone H3 (CitH3), a marker of NETs, were rarely observed except in thin-cap fibroatheromas and ruptured plaques. Fibrin deposition was detected from pathological intimal thickening onward and increased in thin-cap fibroatheromas and ruptured plaques. VWF deposition was mainly observed in necrotic cores and was abundant in ruptured plaques. Among non-ruptured plaques, CitH3- and fibrin-positive areas were significantly larger in plaques with intraplaque hemorrhage than in those without hemorrhage. In cultured human neutrophils, stimulation with hemin, ATP, ADP, or thrombin induced NET formation.

These findings suggest that neutrophils, NETs, and platelets are not prominent features of early human coronary atherosclerotic lesions but become more evident with plaque progression and destabilization. In contrast, fibrin and VWF deposition can already be detected in earlier lesions. Intraplaque hemorrhage may further promote fibrin deposition, and NET-associated responses via erythrocyte-, platelets, coagulation-related molecules. Neutrophil accumulation and VWF deposition may predominantly accompany plaque rupture.

## 1. Introduction

Myocardial infarction is one of the leading causes of death worldwide. Coronary artery atherosclerosis is a gradual process that remains asymptomatic for a long time. However, acute changes in plaques, plaque disruption, and thrombus formation can lead to myocardial infarction. Plaque rupture is a major morphological feature of plaque disruption. Although arterial thrombus formation is initiated by platelet adhesion to the exposed subendothelial collagenous matrix via von Willebrand factor (VWF) [1], the ruptured plaque is characterized by a small amount of subendothelial fibrous matrix. A recent pathological study revealed that fibrin and VWF, rather than a fibrous matrix, are present under platelet aggregation at the ruptured plaque-thrombus interface in patients with acute myocardial infarction [2]. Neutrophil extracellular traps (NETs) are decondensed chromatin and proteins released from neutrophils, which promote thrombus formation [3]. NET formation is also observed at the interface. These findings suggest that fibrin, VWF, and NETs play roles in thrombus formation as scaffolds for platelet adhesion and aggregation in ruptured coronary plaques. However, whether these elements were present before or resulted from the plaque rupture remains unclear.

Previously, neutrophils were not detected in coronary diffuse intimal thickening or fibrous plaques but were detected in 6% of fibrolipid lesions and in all ruptured plaques[4]. Neutrophils and NETs are more abundant in ruptured coronary plaques and plaques with intraplaque hemorrhage (IPH) than in nonthrombosed plaques[5]. Plaque disruption and IPH may associate neutrophilic infiltration and NET formation in the coronary arteries.

Fibrin is rarely observed in early atherosclerotic lesions and is distributed around foam cells in fibrous plaques and necrotic cores in advanced plaques. In advanced lesions, fibrin deposition is detected parallel to the luminal surface, suggesting the incorporation of a mural thrombus [6]. In advanced aortic plaques, VWF was detected together with fibrin and platelet glycoprotein (GP) IIb/IIIa, suggesting intraplaque hemorrhage and/or mural thrombus incorporation [7]. However, these findings for fibrin and VWF are predominantly based on the aorta and peripheral arteries.

According to atherosclerotic lesion progression and thrombotic complications, coronary atherosclerotic lesions are histologically classified as diffuse intimal thickening, pathological intimal thickening, fibrous cap atheroma, fibrocalcified plaque, thin-cap fibroatheroma, and disrupted plaques [8]. Thin-cap fibroatheroma is considered a precursor lesion to ruptured plaque [9]. Although a pathological study reported an increased fibrin deposition score in fibrous cap atheroma and thin-cap fibroatheroma compared with that in diffuse intimal thickening and pathological intimal thickening[10], the presence and distribution of neutrophils, NETs, and VWF remain unknown among the coronary atherosclerosis classification.

In this study, we aimed to reveal the presence and extent of neutrophils, NETs, fibrin, and VWF formation during human coronary atherogenesis and the relationship between these factors and IPH.

## 2. Materials and methods

### 2.1 Study cohort

This study was conducted in accordance with the Declaration of Helsinki. The Ethics Committee of the University of Miyazaki approved the study protocol (approval no. O-1245). As this was a retrospective study of autopsy cases, we applied the opt-out method to obtain consent. Human coronary arteries were obtained from 16 autopsy cases (five cases of non-cardiac death and 11 cases of ischemic heart disease [IHD]). The IHD group included five cases with acute myocardial infarction (Supplementary Table 1). The age of the non-IHD cases tended to be lower than that of the IHD cases, and the IHD cases tended to have a higher frequency of dyslipidemia than non-IHD cases. However, the clinical characteristics of both groups did not significantly differ. Formalin-fixed, paraffin-embedded sections (3 μm thickness) of the coronary arteries (n=1 to 11 in each case) were obtained from the proximal portion of the right and left coronary arteries or the culprit lesion of myocardial infarction and stained with hematoxylin and eosin. We histologically evaluated 102 sections of the human coronary arteries. Coronary lesions were classified according to atherosclerosis classification as diffuse intimal thickening, pathological intimal thickening, fibrous cap atheroma, fibrocalcified plaque, thin-cap (<65 μm) fibroatheroma, or ruptured plaque, as defined by Virmani et al. [8]. We also assessed each lesion for the presence or absence of intraplaque hemorrhage.

### 2.2 Immunohistochemical analysis in human coronary artery

The sections were immunohistochemically analyzed using antibodies against α-SMA (a smooth muscle cell [SMC] marker, mouse monoclonal, 1A4, DAKO/Agilent, Santa Clara, CA, USA), CD68 (a macrophage marker, mouse monoclonal, PG-M1, DAKO/Agilent), CD66b (a neutrophil marker, mouse monoclonal, clone 6/40c; Biolegend Inc., San Diego, California, USA), citrullinated histone H3 (a NET marker, rabbit polyclonal; Abcam Cambridge, Massachusetts, USA), GPII/IIIa (a platelet marker, sheep polyclonal antibody; Affinity Biologicals, Inc., Hamilton, CA, USA), fibrin (mouse monoclonal antibody, clone T2G1; Accurate Chemical & Scientific Corp. Westbury, NY, USA), VWF (mouse monoclonal antibody, clone 36B11, Novocastra, Newcastle upon Tyne, United Kingdom), and glycophorin A (erythrocyte marker; mouse monoclonal antibody, clone JC159; DAKO/Agilent). Immunoreactive signals were visualized using the EnVision System (DAKO). Microscopic digital images of α-SMC, macrophage, GPIIb/IIIa, fibrin, VWF, and glycophorin A immunohistochemistry were captured with a digital camera (DP-74, OLYMPUS, Tokyo, Japan) under a 10× objective lens. Since the immunoreactivity for CD66b and Cit-H3 was focal and subtle in most coronary lesions, microscopic digital images for CD66b and Cit-H3 immunohistochemistry were captured in the most densely stained area in each arterial section using a 40× objective lens. Immunopositive areas for each antigen in the coronary arteries were semi-quantified using a color imaging morphometry system (WinROOF; Mitani, Fukui, Japan) [11]. Briefly, immunopositive areas were extracted as green areas using specific protocols based on the color parameters in the software, namely, hue, lightness, and saturation. Data were expressed as positively stained areas per intimal area in each lesion. Intraplaque hemorrhage was defined as the extravasation of intact erythrocytes in plaques with hematoxylin–eosin stain [12]. Since IPH was observed only in fibrous cap atheroma and thin-cap fibroatheroma among the non-ruptured lesions, these sections were immunohistochemically analyzed using glycophorin A antibody.

### 2.3 Isolation of human peripheral blood neutrophils and in vitro stimulation

The study was approved by the Research Ethics Committee of the University of Miyazaki (approval no. O-2008), and written informed consent was obtained from all healthy volunteers. Human peripheral blood neutrophils were isolated from venous blood by density-gradient centrifugation using Polymorphprep (Serumwerk Bernburg AG, Bernburg, Germany) [13, 14]. The isolated neutrophils were washed with Hanks’ balanced salt solution without calcium, magnesium, or phenol red [HBSS(-); FUJIFILM Wako Pure Chemical Corporation, Osaka, Japan]. Cells were maintained in RPMI-1640 medium supplemented with 10% fetal bovine serum (FBS; Sigma-Aldrich, St. Louis, MO, USA) and 1% ZellShield (Minerva Biolabs, Berlin, Germany) at 37°C in a humidified atmosphere containing 5% CO□. Neutrophils were stimulated with various concentrations of hemin (FUJIFILM Wako Pure Chemical Corporation, Osaka, Japan), ATP (MedChemExpress, Monmouth Junction, NJ, USA), ADP (Toronto Research Chemicals Inc., Toronto, Canada), or thrombin (MP Biomedicals, Irvine, CA, USA), for 1 h. NET-associated responses were subsequently evaluated by immunocytochemistry and immunofluorescence for CitH3.

### 2.4 Immunocytochemistry for CitH3 and quantification in neutrophil culture experiment

After stimulation, the entire cell suspension, including the culture medium, was collected and diluted fivefold. An aliquot (200 μL) of the diluted cell suspension was cytocentrifuged onto glass slides using a Cytospin centrifuge (Cytospin4, PHC corporation, Tokyo, Japan). The cytospin preparations were immediately fixed with 4% paraformaldehyde (PFA) for overnight at 4□.

The slides were incubated with an anti-Histone H3 (citrulline R2 + R8 + R17) antibody (rabbit polyclonal, 1:1,000; ab5103; Abcam, Cambridge, UK). For immunofluorescence, a CF568-conjugated donkey anti-rabbit IgG secondary antibody (1:50, Biotium, Fremont, CA, USA) was used, and slides were mounted with a medium containing 4′,6-diamidino-2-phenylindole (DAPI). Fluorescence images were acquired using a microscope equipped with a DP74 digital camera (Olympus, Tokyo, Japan), and fluorescence channels were merged using CellSens Standard software, version 2.3 (Olympus).

For immunocytochemistry, a goat anti-rabbit IgG secondary antibody (Histofine simplestain MAX-PO(R), Nichirei Biosciences Inc., Tokyo, Japan) was used. Horseradish peroxidase activity was visualized using 3,3’-diaminobenzidine tetrahydrochloride, followed by counterstaining with Mayer’s hematoxylin. For quantitative analysis, 15 randomly selected fields were photographed for each sample using a ×10 objective lens. The percentage of CitH3-positive neutrophils was calculated as the number of CitH3-positive neutrophils divided by the total number of neutrophils in each field.

### 2.5 Statistical analysis

All statistical analyses were performed using the GraphPad Prism 9 software (GraphPad Software Inc., San Diego, CA, USA). The Shapiro-Wilk test was used for normality analysis. Data are expressed as medians and interquartile ranges, or as individual data points. Variables between or among groups were compared using Mann-Whitney U tests or Kruskal-Wallis tests, followed by Dunn’s multiple comparison test. Contingency analysis was performed using Fisher’s exact test. Correlations between factors were evaluated using Spearman’s test. P-values of <0.05 were considered statistically significant.

## 3. Results

### 3.1 Distribution of coronary atherosclerotic plaque types in autopsy cases with and without ischemic heart disease

We histologically examined 102 coronary artery sections from five non-IHD cases (37 sections) and 11 IHD cases (65 sections). Coronary lesions were classified as diffuse intimal thickening, pathological intimal thickening, fibrous cap atheroma, fibrocalcified plaque, thin-cap fibroatheroma, or ruptured plaque according to the established classification of atherosclerotic lesions [8]. Figure 1A shows the distribution of these plaque types in IHD and non-IHD cases. In non-IHD cases, diffuse intimal thickening was the most common lesion (55%), followed by pathological intimal thickening (31%). Fibrous cap atheroma, thin-cap fibroatheroma, and ruptured plaque were not observed in non-IHD cases. In contrast, IHD cases exhibited a broader spectrum of atherosclerotic lesions, including pathological intimal thickening, fibrous cap atheroma, fibrocalcified plaque, thin-cap fibroatheroma, and ruptured plaque. Diffuse intimal thickening was the least common lesion in IHD cases in contrast to the non-IHD cases.

**Fig. 1.**
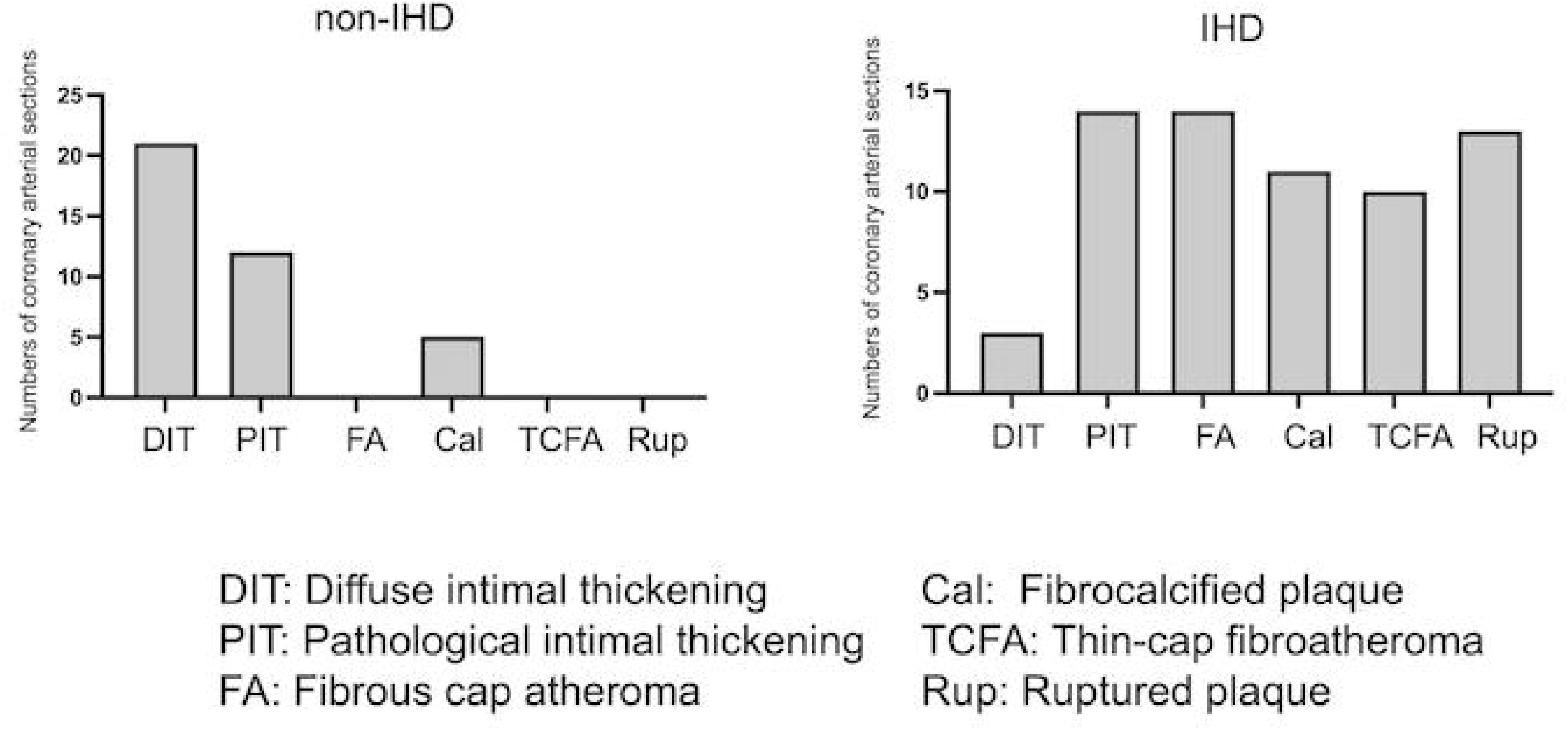
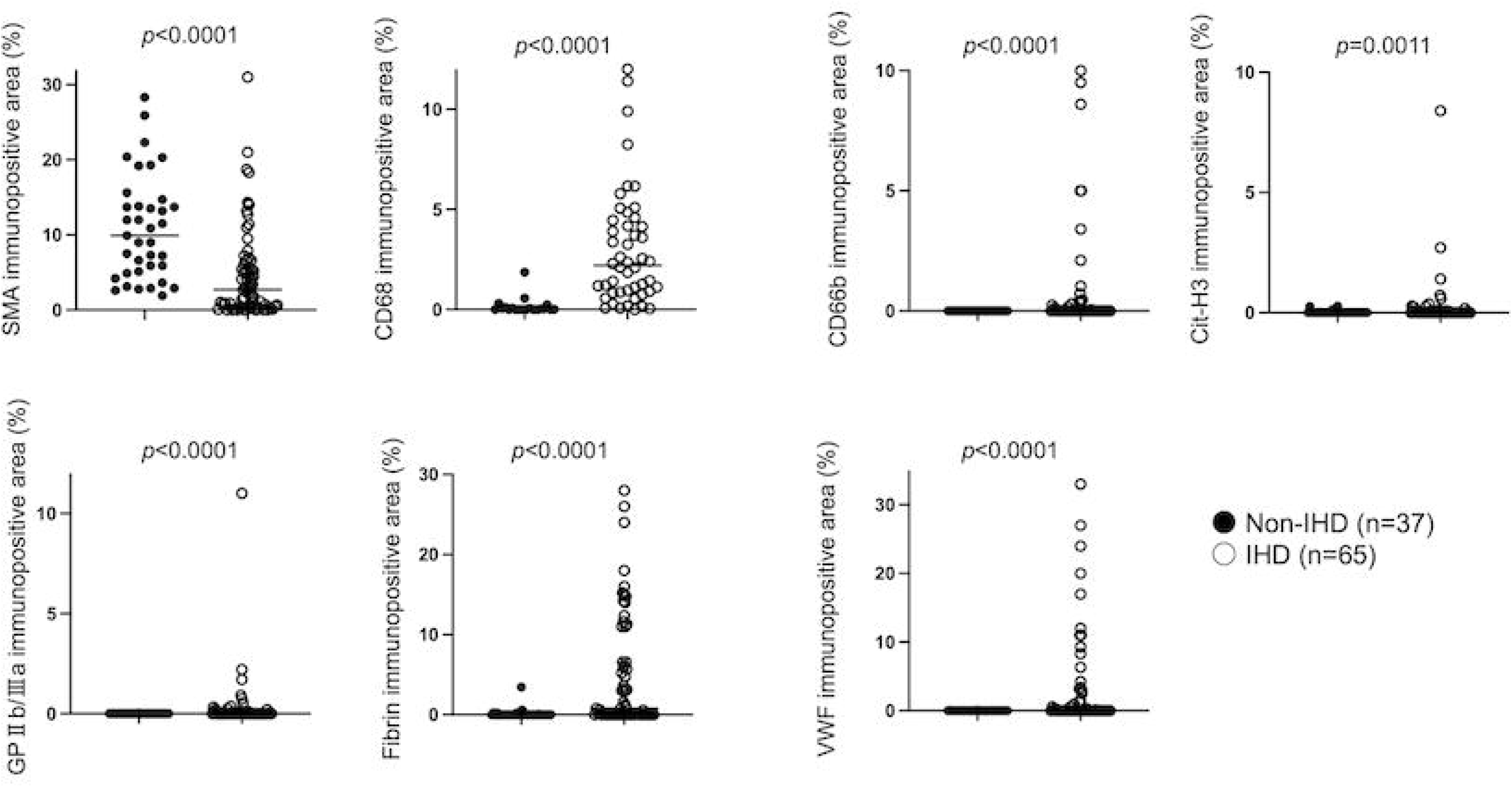
Distribution of atherosclerotic plaque classification and immunopositive area for cellular and molecular components in coronary arteries in autopsy cases with or without ischemic heart disease (A) Distribution of atherosclerotic plaque classification in the human coronary artery in autopsy cases with or without ischemic heart disease (IHD). (B) Immunopositive areas for α-smooth muscle actin (SMA), CD68, CD66b, and citrullinated histone H3 (CitH3), glycoprotein (GP)IIb/IIIa, fibrin, and von Willebrand factor (VWF) in human coronary arteries in autopsy IHD and non-IHD cases. Data were analyzed using Mann-Whitney U tests

### 3.2 Distribution of neutrophils, NETs, platelets, fibrin, and VWF in coronary atherosclerotic lesions

Immunohistochemistry was performed for SMA, CD68, CD66b, CitH3, GPIIb/IIIa, fibrin, and VWF to evaluate smooth muscle cells, macrophages, neutrophils, NET-associated changes, platelets, and thrombotic components. Compared with non-IHD cases, coronary arteries from IHD cases showed greater immunopositive areas for CD68, CD66b, CitH3, GPIIb/IIIa, fibrin, and VWF (Fig. 1B).

Representative histological and immunohistochemical findings according to plaque type are shown in Fig. 2. Diffuse intimal thickening was characterized by abundant SMA-positive smooth muscle cells and sparse macrophages, with little or no CD66b, CitH3, GPIIb/IIIa, fibrin, or intimal VWF staining (Fig. 2A). Pathological intimal thickening showed focal macrophage infiltration and slight fibrin and VWF deposition, whereas CD66b, CitH3, and GPIIb/IIIa showed little to no staining (Fig. 2B). Fibrous cap atheromas contained macrophage-rich necrotic cores with mild fibrin and VWF deposition but little CD66b, CitH3, or GPIIb/IIIa immunoreactivity (Fig. 2C). Fibrocalcified plaques showed extensive calcification with sparse macrophages and few or no staining for CD66b, CitH3, GPIIb/IIIa, fibrin, and VWF (Fig. 2D).Thin-cap fibroatheromas were characterized by a thin fibrous cap containing few SMCs and a large necrotic core with cholesterol clefts, with or without intraplaque hemorrhage. They showed marked macrophage infiltration, increased cellular and extracellular CitH3 and CD66b immunoreactivity, and greater fibrin and VWF deposition within the necrotic core; GPIIb/IIIa-positive platelets were also present in some lesions (Fig. 2E). Ruptured plaques showed plaque disruption with platelet-fibrin thrombus formation, dense macrophage and neutrophil infiltration, prominent CitH3 immunoreactivity, and abundant fibrin, VWF, and platelet deposition (Fig. 2F). Mural platelet-fibrin thrombi were not observed in non-ruptured plaque types.

**Fig. 2.**
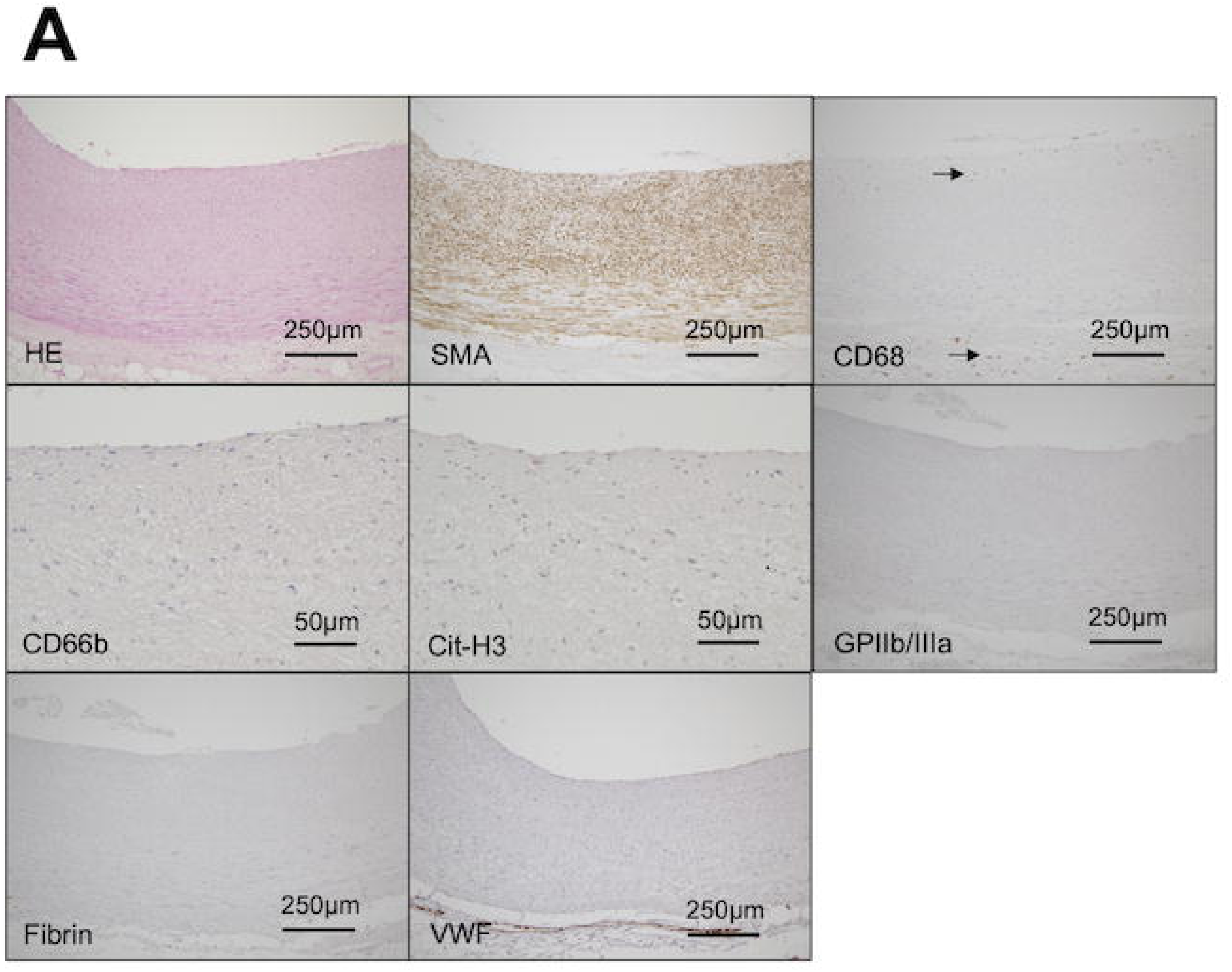
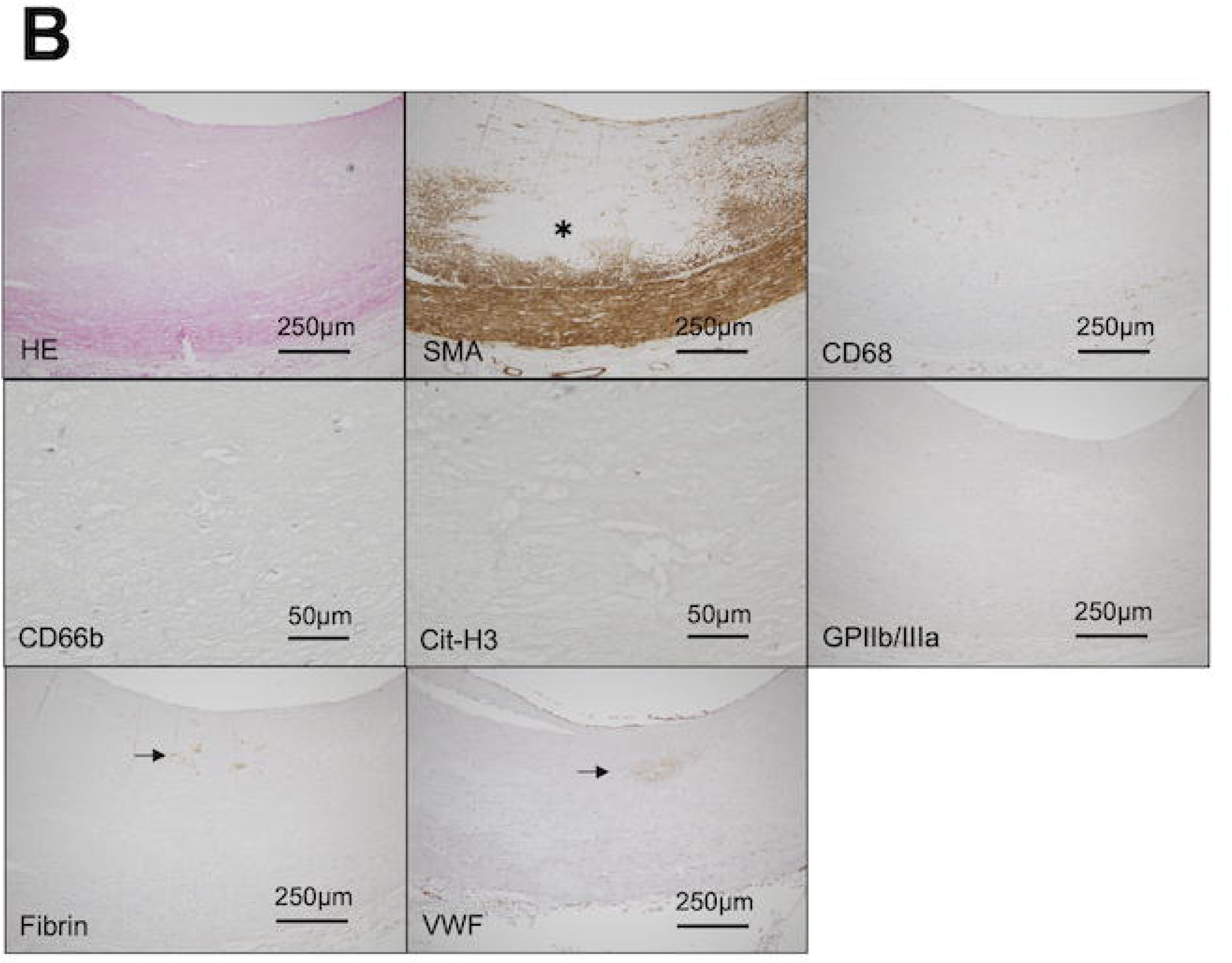
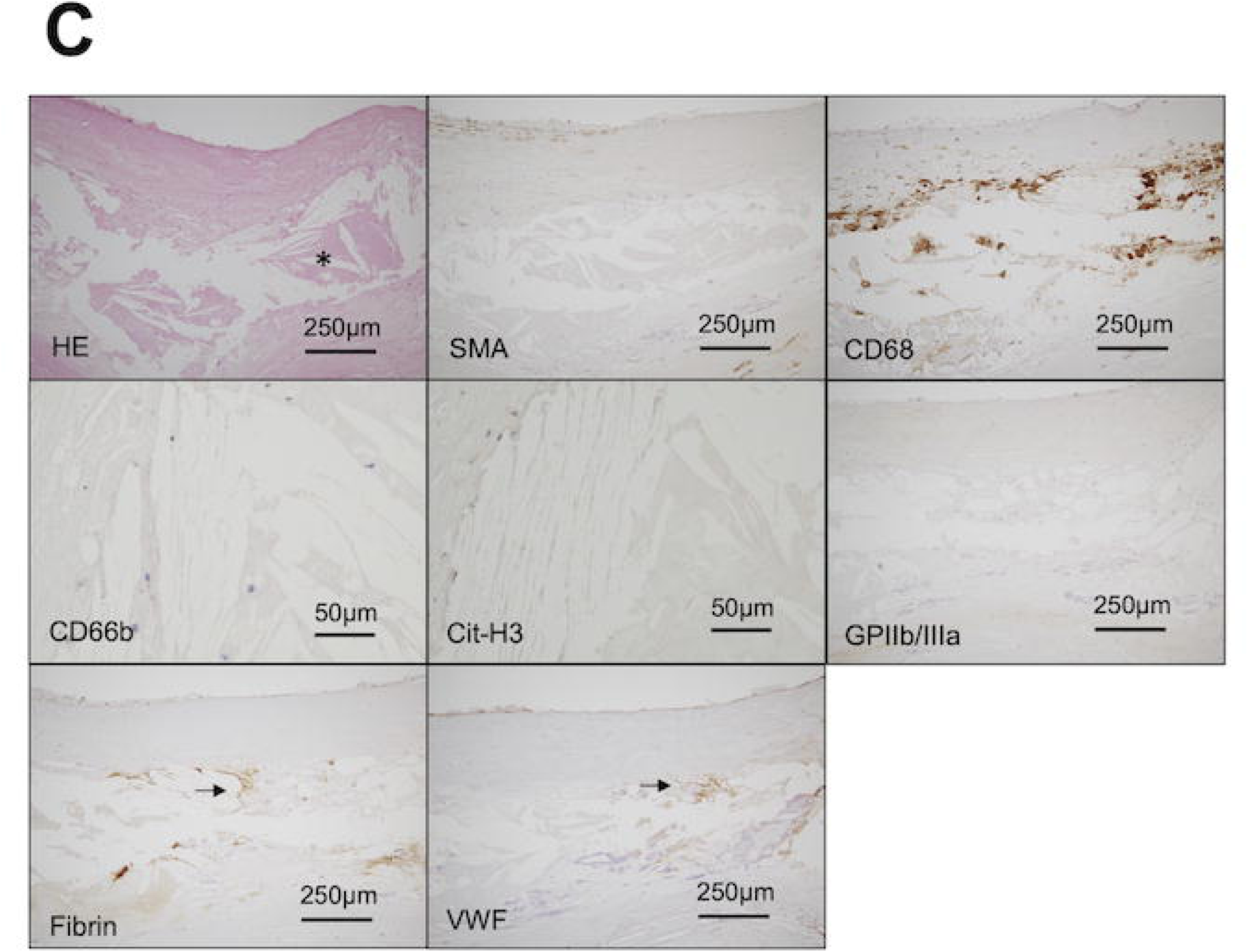
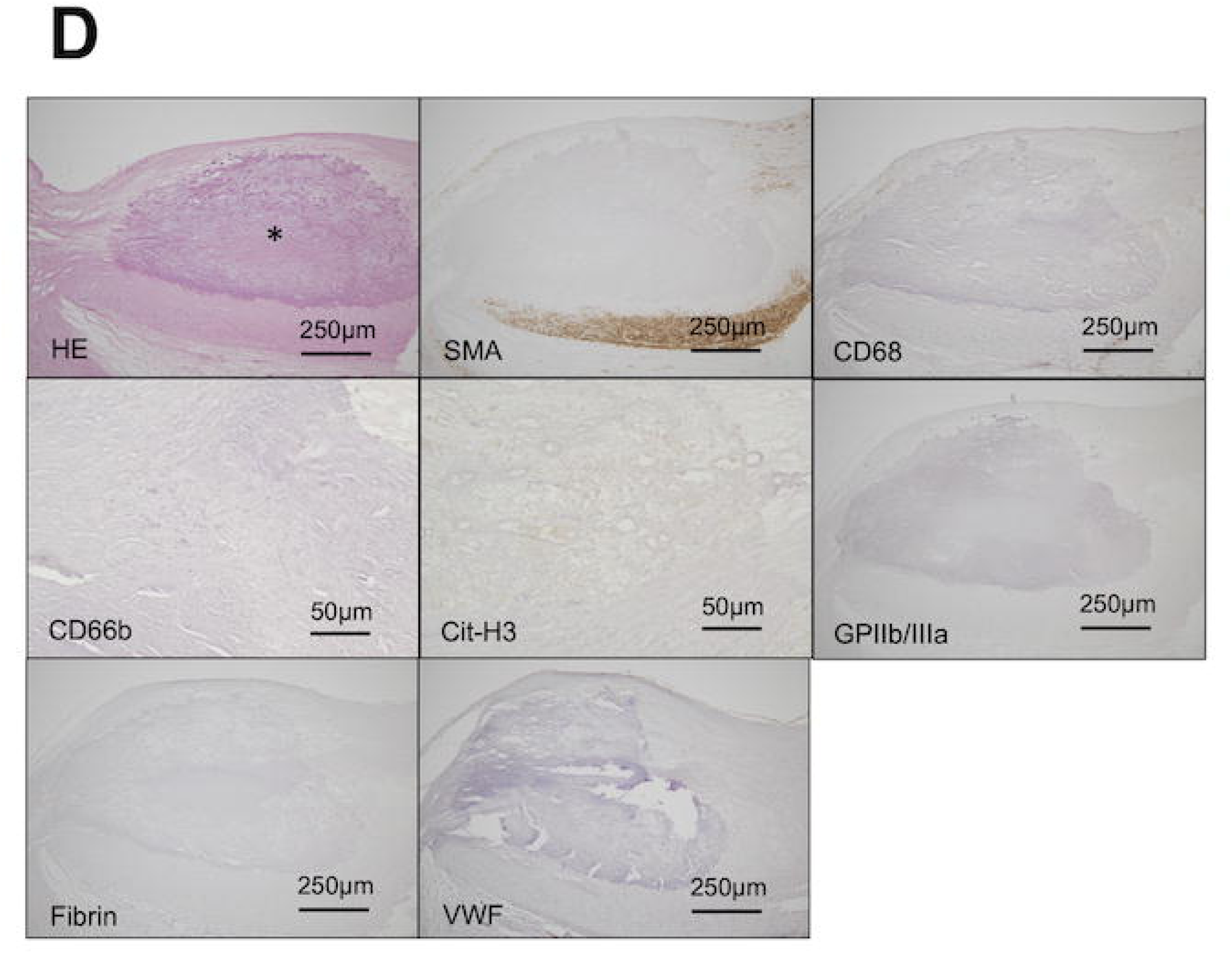
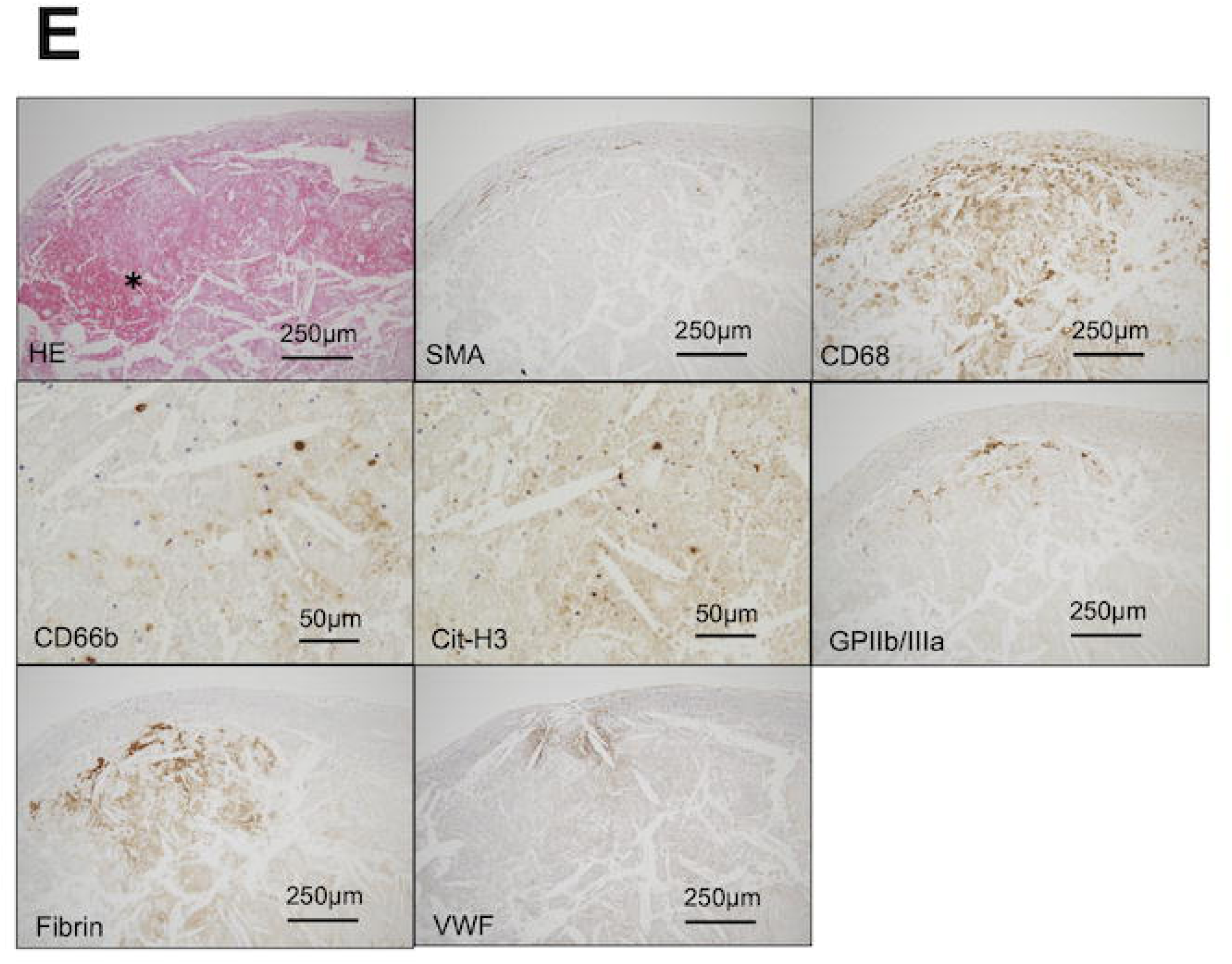
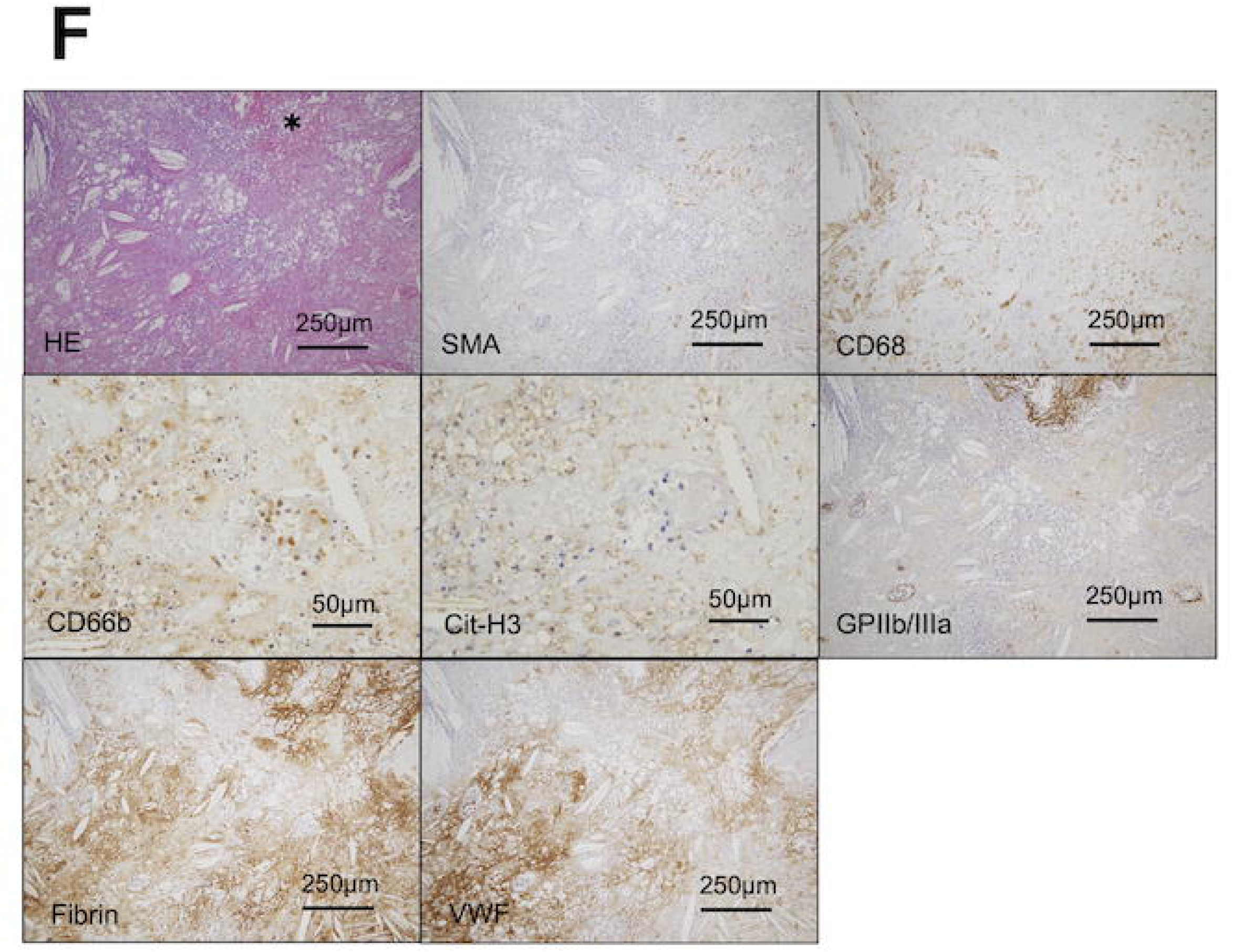
Representative histological images of each classification of human coronary atherosclerosis and expressions of SMA, CD68, CD66b, and CitH3, GPIIb/IIIa, fibrin, and VWF. (A) Representative histological and immunohistochemical images of diffuse intimal thickening (DIT). DIT exhibits a diffuse distribution of SMA-positive smooth muscle cells (SMC) in the intima and media, and a sparse distribution of CD68 macrophages in the intima and adventitia (arrows). DIT is immunonegative for CD66b, CitH3, GPIIb/IIIa, fibrin, and VWF in the intima, except for a luminal immunoreaction for endothelial VWF. (B) Representative histological and immunohistochemical images of pathological intimal thickening (PIT). PIT exhibits partial loss of SMCs with the accumulation of extracellular matrix or lipids (asterisk) in the intima. PIT shows slight infiltration of macrophages and slight fibrin and VWF deposition (arrows) in the intima. PIT is immunonegative for CD66b, CitH3, and GPIIb/IIIa in the intima. (C) Representative histological and immunohistochemical images of the fibrous cap atheroma. The fibrous cap atheroma has a necrotic core with cholesterol clefts (asterisk) and a fibrous cap. The fibrous cap atheroma exhibits SMCs in the fibrous cap, infiltration of macrophages in the necrotic core, and mild deposition of fibrin and VWF in the necrotic core (arrows). Fibrous cap atheromas are immunonegative for CD66b, CitH3, and GPIIb/IIIa. (D) Representative histological and immunohistochemical images of fibrocalcified plaques. The fibrocalcified plaque exhibits broad calcification (asterisk), focal presence of SMCs, and sparse distribution of macrophages in the plaque. The fibrocalcified plaque was immunonegative for CD66, CitH3, GPIIb/IIIa, fibrin, and VWF. (E) Representative histological and immunohistochemical images of thin-cap fibrous atheroma (TCFA). TCFA has a thin fibrous cap and a large necrotic core with intraplaque hemorrhage (asterisk) and cholesterol clefts. TCFA induces infiltration by large numbers of macrophages, cellular or extracellular immunoreactivity for CD66b and CitH3, and the deposition of fibrin and VWF in the necrotic core. GPIIb/IIIa-positive platelets were observed under a thin fibrous cap. (F) Representative histological and immunohistochemical images of ruptured plaques. The ruptured plaque exhibited disruption of the atherosclerotic plaque and platelet-fibrin thrombus formation (asterisk), infiltration of large numbers of CD68-positive macrophages, dense cellular or extracellular immunoreactivity for CD66b and CitH3, presence of platelets, and deposition of fibrin and VWF in the disrupted plaque HE, hematoxylin-eosin, Scale bars represent 250 μm or 50 μm

Quantitative data are summarized in Supplementary Table 2. CD66b, CitH3, and fibrin were increased in thin-cap fibroatheromas and ruptured plaques, whereas GPIIb/IIIa and VWF were most prominent in ruptured plaques.

### 3.3 Expression of CD66b, CitH3, fibrin, and VWF in human coronary stable plaques

To examine the cellular components, NETs formation, deposition of fibrin, and VWF in stable plaques, pathological intimal thickening, fibrous cap atheroma, and fibrocalcified plaques, we compared the immunopositive areas for SMA, CD68, CD66b, Cit-H3, GPIIb/IIIa, fibrin, and VWF (Figure 3). The area immunopositive for SMA in pathological intimal thickening was greater than that in fibrous cap atheromas and fibrocalcified plaques. Fibrous cap atheroma had a higher level of macrophage and neutrophil infiltration, as well as fibrin deposition, than those in pathological intimal thickening and fibrocalcified plaques. VWF deposition in fibrous cap atheromas is greater than that in pathological intimal thickening. No significant differences were observed in NET formation or GP IIb/IIIa expression among the groups.

**Fig. 3.**
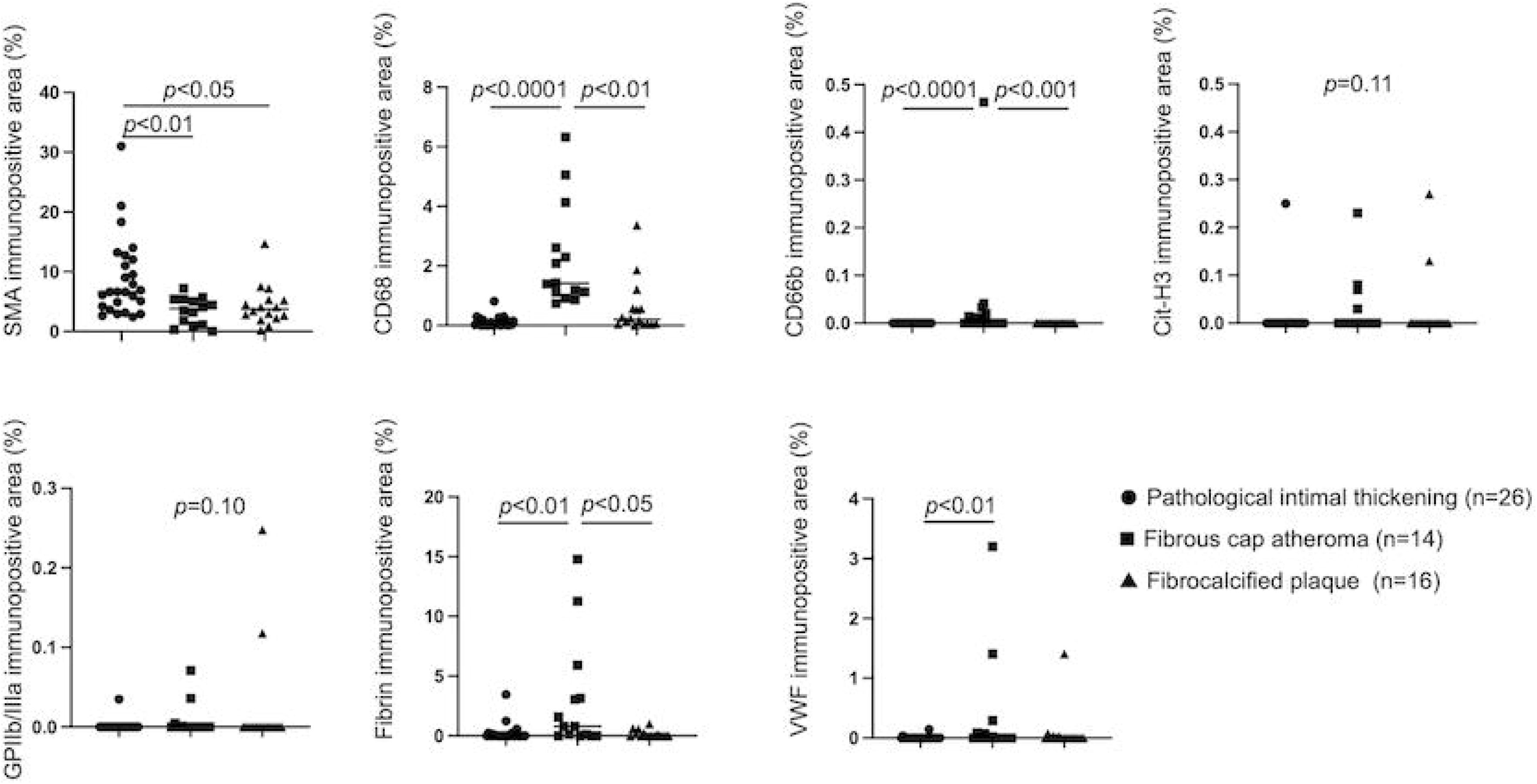
Expression of SMA, CD68, CD66b, CitH3, GPIIb/IIIa, fibrin, and VWF in coronary stable lesions. Immunopositive areas for α-smooth muscle actin (SMA), CD68, CD66b, citrullinated histone H3 (CitH3), glycoprotein (GP) IIb/IIIa, fibrin, and von Willebrand factor (VWF) in pathological intimal thickening, fibrous cap atheroma, and fibrocalcified plaques in human coronary arteries. Data were analyzed using Kruskal-Wallis tests, followed by Dunn’s multiple comparison tests

### 3.4 Expression of cellular and thrombotic markers across atheromatous coronary plaque types

To compare cellular components and thrombotic markers among fibrous cap atheromas, thin-cap fibroatheromas, and ruptured plaques, we quantified the immunopositive areas for SMA, CD68, CD66b, CitH3, GPIIb/IIIa, fibrin, and VWF (Fig. 4). The SMA-positive area was greater in fibrous cap atheromas than in thin-cap fibroatheromas and ruptured plaques. In contrast, macrophage infiltration and the CitH3- and fibrin-positive areas were greater in thin-cap fibroatheromas and ruptured plaques than in fibrous cap atheromas. Neutrophil infiltration, platelet accumulation assessed by GPIIb/IIIa, and VWF deposition were more prominent in ruptured plaques compared with fibrous cap atheromas and thin-cap fibroatheromas.

**Fig. 4.**
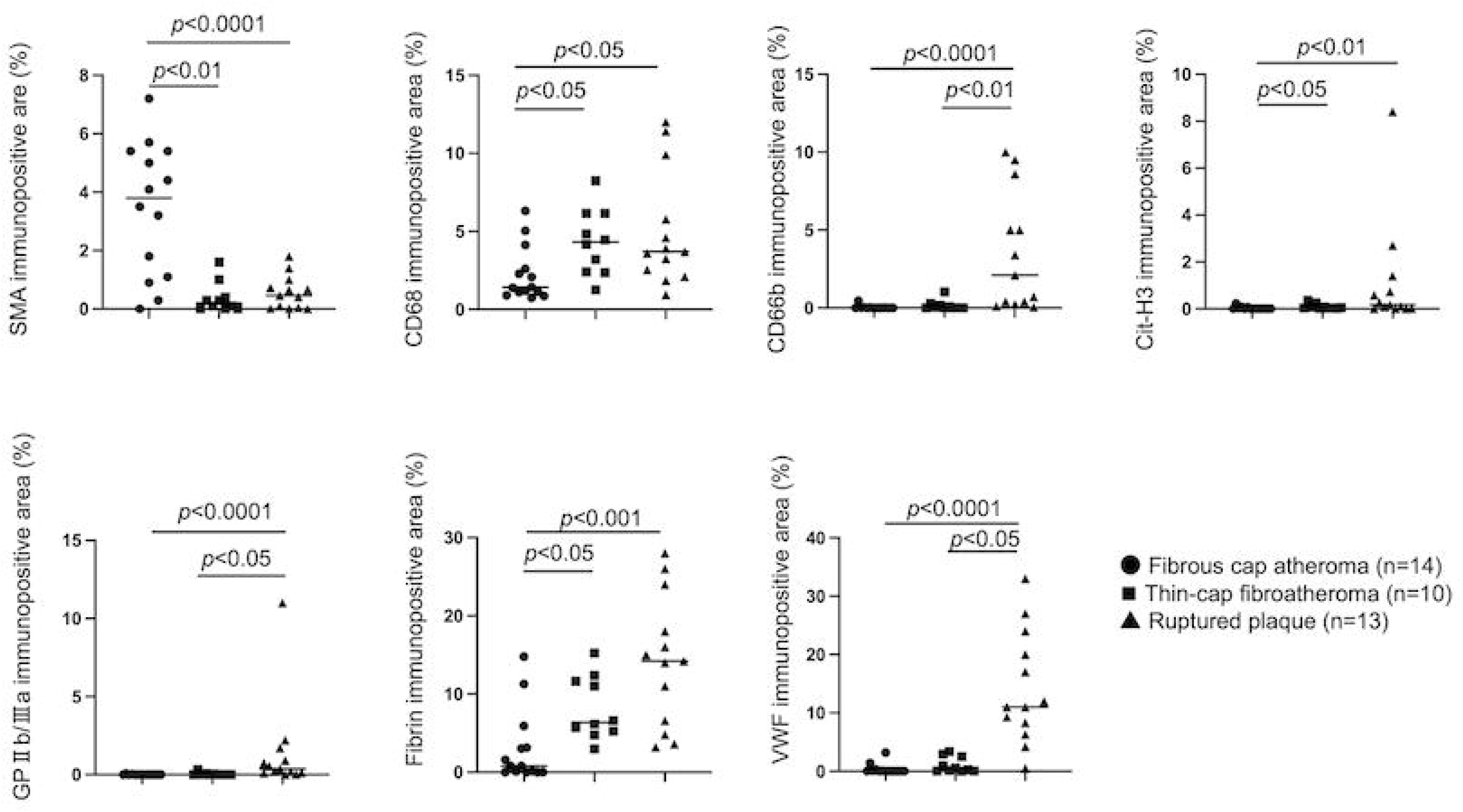
Expression of SMA, CD68, CD66b, CitH3, GPIIb/IIIa, fibrin, and VWF in coronary atheromas. Immunopositive areas for α-smooth muscle actin (SMA), CD68, CD66b, citrullinated histone H3 (CitH3), glycoprotein (GP)IIb/IIIa, fibrin, and von Willebrand factor (VWF) in fibrous cap atheroma, thin-cap fibrous atheroma, and ruptured plaques. Data were analyzed using Kruskal-Wallis tests, followed by Dunn’s multiple comparison tests

### 3.5 Association of intraplaque hemorrhage with plaque components

Intraplaque hemorrhage (IPH) was observed in 35% of fibrous cap atheromas and 90% of thin-cap fibroatheromas among non-ruptured lesions. IPH was not observed in diffuse intimal thickening, pathological intimal thickening, or fibrocalcified plaques. Ruptured plaques were excluded from this analysis because hemorrhage associated with plaque rupture could confound the assessment of spontaneous IPH.

Figure 5 shows the immunopositive areas for SMA, CD68, CD66b, CitH3, GPIIb/IIIa, fibrin, and VWF in fibrous cap atheromas and thin-cap fibroatheromas with or without IPH. The SMA-positive area was significantly smaller in atheromas with IPH than in those without IPH. In contrast, CitH3-positive area, platelet accumulation, and fibrin deposition were significantly greater in atheromas with IPH. No significant differences were observed in macrophage or neutrophil infiltration or VWF deposition according to the presence or absence of IPH.

**Fig. 5.**
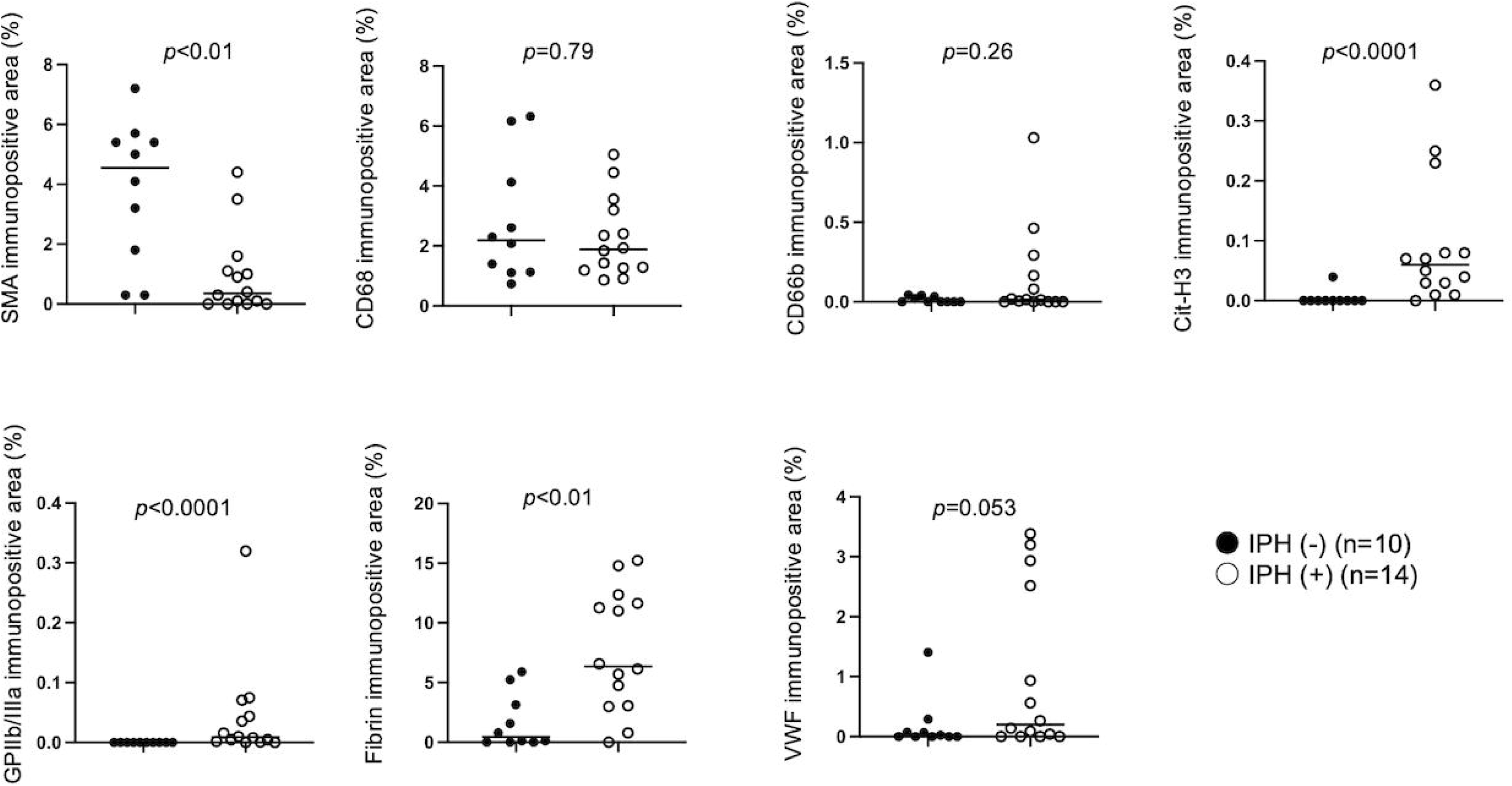
Expression of SMA, CD68, CD66b, CitH3, GPIIb/IIIa, fibrin, and VWF in coronary plaques with or without intraplaque hemorrhage. Immunopositive areas for α-smooth muscle actin (SMA), CD68, CD66b, and citrullinated histone H3 (CitH3), glycoprotein (GP)IIb/IIIa, fibrin, and von Willebrand factor (VWF) in coronary plaques with or without intraplaque hemorrhage (IPH). Data were analyzed using the Mann-Whitney U test

To quantify erythrocyte accumulation within plaques, immunohistochemistry for glycophorin A, an erythrocyte marker, was performed. The glycophorin A-positive area was significantly greater in thin-cap fibroatheromas (median [interquartile range], 2.2% [0.7–5.0%], n = 10) than in fibrous cap atheromas (0% [0–0.35%], n = 14; p < 0.01). The glycophorin A-positive area positively correlated with the CitH3-positive area (r = 0.73, p < 0.0001, n = 24) and the GPIIb/III-positive area (r=0.80, *p*<0.0001, n=24), fibrin-positive area (r=0.75, *p*<0.0001, n=24), and VWF-positive area (r=0.48, *p*=0.015, n=24).

### 3.6 Effects of hemin, ATP, ADP, and thrombin on NET-associated responses in human peripheral blood derived neutrophils

To evaluate whether IPH can induce NETs formation in neutrophils, peripheral blood derived neutrophils were stimulated with erythrocyte-, platelet-, coagulation-related molecules; hemin, ATP, ADP, and thrombin. After stimulation of cultured neutrophils with 20 μM hemin, 25 μM ATP, 25 μM ADP, or 1 U/mL thrombin for 1 h, immunofluorescence demonstrated filamentous DNA co-localized with CitH3 (Fig. 6A). Representative images of CitH3 immunocytochemistry showed filamentous CitH3-positive material around neutrophils stimulated with 20 μM hemin, 25 μM ATP, 25 μM ADP, or 1 U/mL thrombin, whereas such staining was less prominent under control conditions (Fig. 6B). The proportion of CitH3-positive neutrophils was significantly increased after stimulation with hemin (10, or 20 μM), ATP (5, or 25 μM), ADP (5, or 25 μM), and thrombin (0.5, or 1 U/mL) compared with the control group (Fig. 6C).

**Fig. 6.**
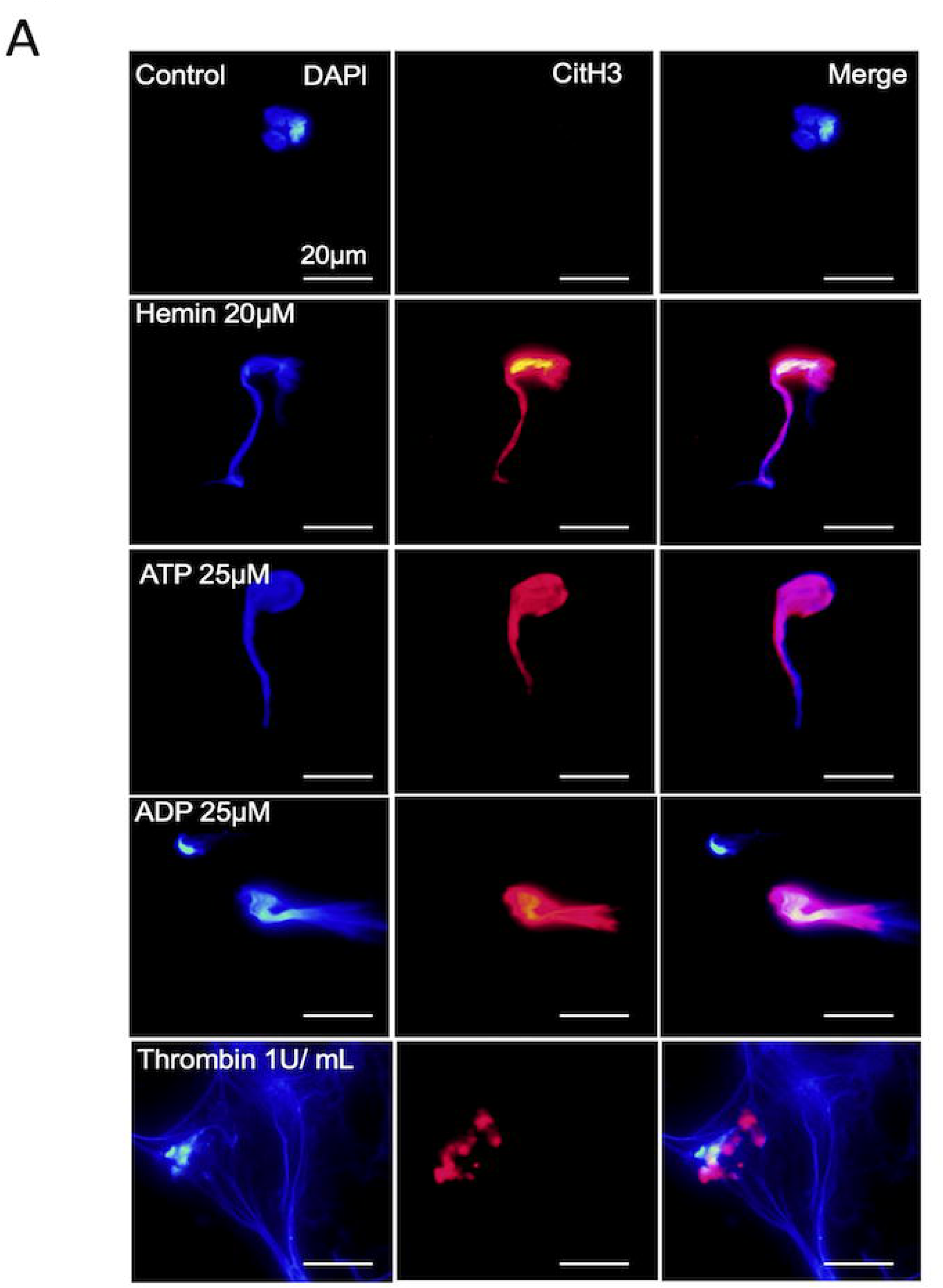
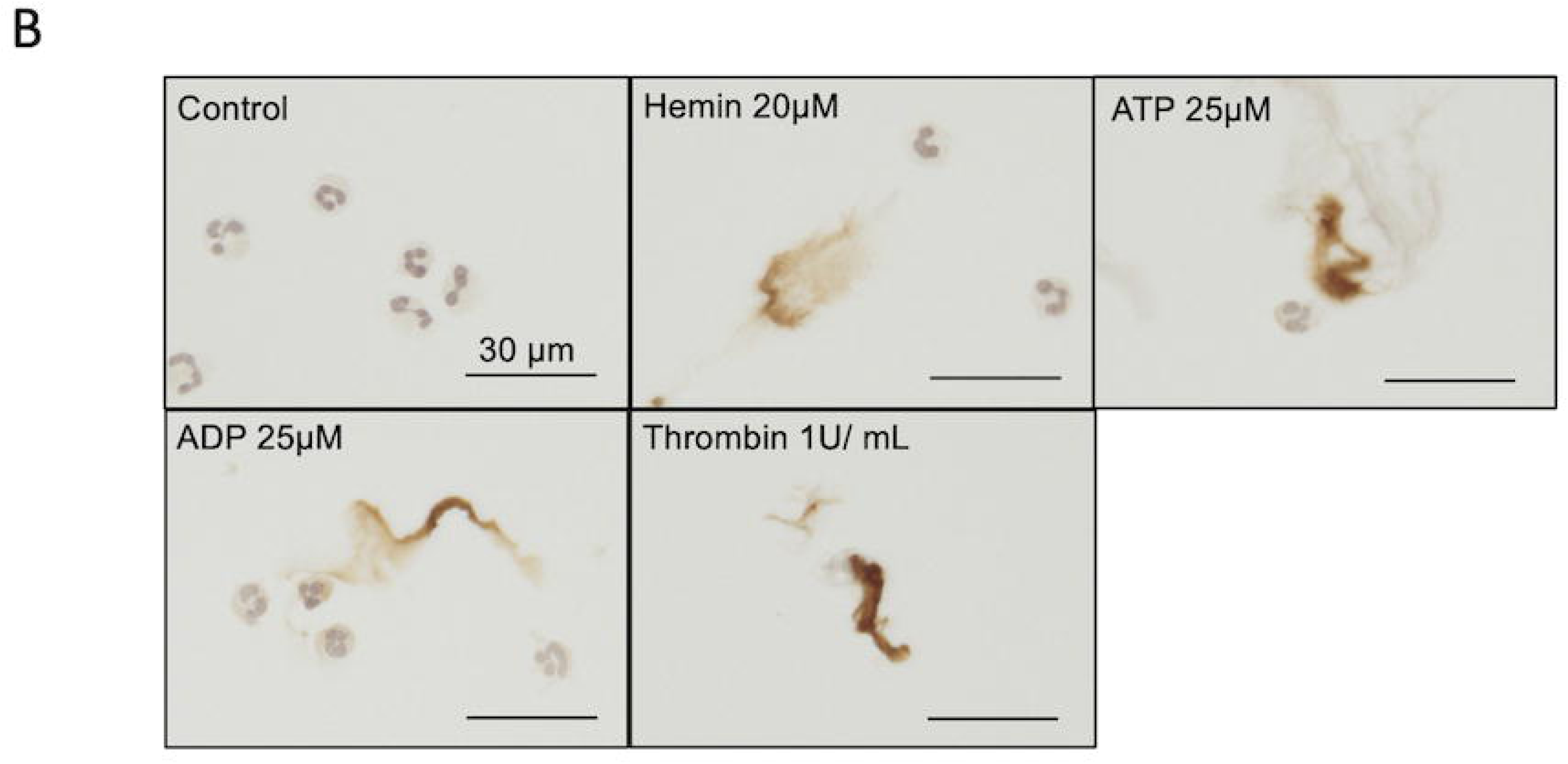
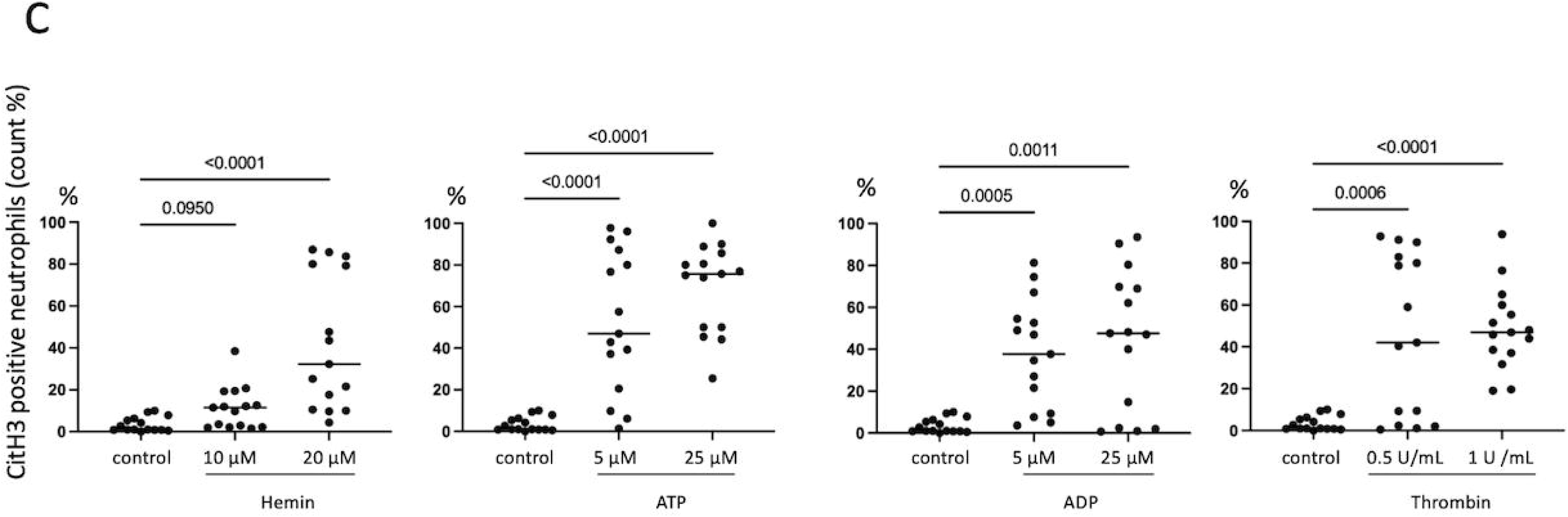
Effects of hemin, ATP, ADP, or thrombin on NET formation in human peripheral blood derived neutrophils. (A) Representative immunofluorescence images showing co-localization of CitH3 (red) with filamentous DNA (blue) in cultured human neutrophils after stimulation with hemin, ATP, ADP, or thrombin. (B) Representative immunocytochemical images of filamentous CitH3 staining in cultured human neutrophils after stimulation with hemin, ATP, ADP, or thrombin. (C) Percentage of CitH3-positive neutrophils after stimulation with hemin, ATP, ADP, or thrombin. Data were analyzed using the Kruskal–Wallis test followed by Dunn’s multiple-comparison test.

## 4. Discussion

In this study, we assessed the distribution and extent of SMCs, macrophages, neutrophils, NETs, platelets, fibrin, and VWF formation during human coronary atherosclerosis using coronary sections from autopsy cases. This study showed that neutrophils, CitH3 expression, and platelets were absent or rare in the coronary plaques of non-IHD cases, and in coronary lesions except for thin-cap fibroatheroma and ruptured plaques. Fibrin and VWF deposition were observed in the pathological intimal thickening and necrotic core of the fibrous cap atheroma. Fibrin deposition was abundant in thin-cap fibroatheromas and ruptured plaques, whereas VWF was abundant in the ruptured plaques. In non-ruptured plaques, the immunopositive areas for CitH3, platelets, and fibrin were larger in plaques with IPH than in non-IPH. In cultured human neutrophils, stimulation with hemin, ATP, ADP, or thrombin promoted NETs formation.

### 4.1 Association of NETs with coronary plaque progression and instability

Neutrophils generate threads of chromatin covered with granule-derived enzymes, namely NETs, in response to injury. A pathological study using human coronary arteries showed that CitH3 immunoreactivity was abundant in all types of complicated plaques, including IPH, eroded plaques, and ruptured plaques, but less frequently in non-thrombosed plaques [5]. Our findings are comparable with those of a previous study. Acute plaque changes, including IPH and plaque disruption, can induce NET formation. In addition, we observed increased CitH3 expression in thin-cap fibroatheromas, a precursor lesion of ruptured plaques. CitH3 was predominantly expressed around the cholesterol clefts in the macrophage-rich necrotic core. Warnatsch et al. reported that cholesterol crystals induce NET formation in human blood-derived neutrophils, and that cholesterol crystals and NETs promote cytokine production in human blood-derived monocytes [13]. Therefore, cholesterol crystal-induced NETs may destabilize thin-cap fibroatheromas through macrophage activation. ApoE-deficient mice show enhanced NET formation in plaques, and inhibition of NET formation protects against atherosclerotic plaque formation [15]. However, in the present study, CitH3 expression was absent or rare in the coronary plaques of cases with non-IHD and coronary plaques, except for thin-cap fibroatheroma and ruptured plaques. Naruko et al. reported that neutrophils were absent or rare in diffuse intimal thickening, fibrous plaques, and fibrolipid plaques [4]. This pathological evidence suggest that NETs play little role in diffuse and pathological intimal thickening, fibrous cap atheroma without IPH, or fibrocalcified atheroma in human coronary arteries.

### 4.2 Potential mechanisms linking intraplaque hemorrhage to NET formation

CitH3-positive NETs, platelet-, fibrin, and VWF depositions were significantly more abundant in human atherosclerotic plaques with intraplaque hemorrhage (IPH) than in those without IPH. To explore possible mechanisms linking IPH to NET formation, we examined the effects of erythrocyte-, platelet-, and coagulation-related molecules that may be present in the hemorrhagic microenvironment. Hemin is a heme-related product generated following erythrocyte breakdown, ATP may be released from damaged blood cells. ADP is released by platelets, and thrombin is generated in blood coagulation. Previous studies demonstrated that hemin and thrombin-activated platelets induces NET formation in human neutrophils [16,17]. These observations, together with our findings, suggest that intraplaque hemorrhage leads to the release of erythrocyte-, platelet-, and coagulation-related molecules, which in turn promote NET formation. Thrombin alone had little effect on NET formation [17]. This may be due to difference in thrombin concentration; 0.5 and 1.0 U/mL in the present study and lower level of thrombin (0.05U/mL) in the previous study.

### 4.3 Distribution of fibrin deposition across plaque types and intraplaque hemorrhage

Fibrin deposition was observed in pathological intimal thickening and in the necrotic cores of atheromas and was particularly abundant in thin-cap fibroatheromas and ruptured plaques. Bini et al. examined fibrin deposition in early and advanced atherosclerotic lesions and reported that fibrin was absent or only minimally present in early lesions but was consistently detected in the necrotic cores of advanced aortic plaques [6]. These findings are consistent with our observations in human coronary arteries. Previous studies have suggested that plaque angiogenesis, rather than IPH itself, is associated with intimal fibrin deposition [10]. Increased permeability of intraplaque microvessels may promote plasma leakage and fibrin deposition even in non-hemorrhagic lesions. In contrast, IPH may further enhance fibrin accumulation in advanced plaques, including fibrous cap atheromas and thin-cap fibroatheromas.

### 4.4 Neutrophilic infiltration associated with plaque disruption

The distribution of neutrophilic infiltration broadly paralleled that of CitH3 expression across coronary plaque types, although the two were not completely concordant. Neutrophilic infiltration was significantly greater in ruptured plaques than in thin-cap fibroatheromas. Previous pathological studies have similarly demonstrated increased neutrophil accumulation in ruptured and eroded coronary plaques [4,5]. These findings suggest that marked neutrophilic infiltration is associated primarily with acute plaque disruption and subsequent thrombus formation.

### 4.5 VWF deposition in relation to plaque progression and intraplaque hemorrhage

VWF deposition was significantly greater in ruptured plaques than in thin-cap fibroatheromas, suggesting that marked VWF accumulation is closely associated with plaque rupture and subsequent thrombus formation. In non-ruptured plaques, the positive association between erythrocyte content and VWF/platelet deposition suggests that IPH may contribute to the accumulation of VWF and platelets within the plaque.

However, the distribution and extent of VWF deposition did not completely correspond to those of platelets. The presence of VWF deposition without platelet accumulation in non-hemorrhagic fibrous cap atheromas suggests that plasma-derived VWF may also enter and be retained within the plaque independently of IPH. Experimental studies have shown that VWF promotes platelet and monocyte adhesion and enhances atherosclerotic lesion formation in LDL receptor-deficient mice [18]. Similarly, subendothelial VWF in the neointima promotes platelet adhesion in cholesterol-fed rabbits [19]. These findings suggest that VWF may contribute to atherogenesis by facilitating the recruitment of platelets and monocytes. In the present human coronary specimens, however, VWF deposition was absent or scant in diffuse intimal thickening and pathological intimal thickening, and mural platelet thrombi were not observed in non-ruptured lesions. Thus, VWF-mediated platelet recruitment demonstrated in experimental models may represent a transient process rather than a persistent feature throughout human coronary atherogenesis.

## 5. Limitations of the study

This study had several limitations. First, because our findings were primarily based on histopathological observations, the causal roles of neutrophils, NETs, fibrin, and VWF in coronary atherogenesis could not be fully established. Although in vitro experiments provided mechanistic insights into NET formation, further studies are needed to clarify the underlying mechanisms. Second, measuring the immunopositive area is a semi-quantitative method that may not necessarily reflect the amount of these factors. However, the fact that neutrophils, NETs, and platelets are virtually absent in lesions other than thin-cap fibroatheromas and ruptured plaques provides important information regarding their involvement in atherogenesis.

## 6. Conclusions

Our histopathological examination of human coronary atherosclerosis suggests that NET formation is rare in stable coronary plaques and that fibrin deposition begins at the stage of pathological intimal thickening and increases with plaque destabilization. IPH may contribute fibrin deposition and NET formation through erythrocyte-, platelet-, and coagulation-related molecules in non-ruptured atheromas. The abundance of neutrophilic infiltrates and VWF deposition in ruptured plaques may reflect a response to plaque rupture.

## Acknowledgments

We thank Nahoko Udatsu and Kyoko Ohashi for their technical assistance in this study. This study was supported by a Clinical Research Support Grant from our institution.

This project was supported by Grants-in-Aid for Scientific Research or Early Career Scientists JSPS KAKENHI (Grant Nos. 19K16560, 19H03445, 23K19470, and 24K18404).

## CRediT authorship contribution statement

EN: Conceptualization, Methodology, Investigation, Formal analysis, Visualization, Writing – original draft. SH: Investigation, Formal analysis. KM: Investigation, review & editing. EK: Methodology, Investigation.TG: Investigation, review & editing. NO: Investigation, review & editing. AM: Supervision, review & editing. MGS: Supervision, review & editing. YA: Supervision, review & editing. AY: Conceptualization, Methodology, Supervision, Writing - review & editing.

## Data availability statement

The datasets used and/or analyzed during the current study are available from the corresponding authors upon reasonable request.

## Ethics approval statement

The Ethics Committee of the University of Miyazaki approved the autopsy study protocol (approval no. O-1245). As this was a retrospective study of autopsy cases, an opt-out approach was used to obtain consent. All data were anonymized. The study involving human peripheral blood was also approved by the Research Ethics Committee of the University of Miyazaki (approval no. O-2008), and written informed consent was obtained from all healthy volunteers.

## Supplementary material

Supplementary Table 1

Supplementary Table 2

**Graphic abstract**

