## Supplemental Table 1 for "Neutrophil extracellular traps and thrombotic components across human coronary plaque progression: Association of intraplaque hemorrhage with NET formation"

**Supplementary table 1. Clinical background of autopsy cases**

| | non-IHD (n=5) | IHD (n=11) | *p* value |
|---|---|---|---|
| Age (years) (median, interquartile range) | 67 (64-72) | 75.5 (68.75-77.75) | 0.15 |
| Male sex | 3 (60%) | 8 (72%) | >0.99 |
| Smoking | 2 (40%) | 5 (45%) | >0.99 |
| BMI (median, interquartile range) | 22.9 (17.05-26.65) | 23.35 (22.9-26.88) | 0.51 |
| Diabates | 1 (20%) | 2 (18%) | >0.99 |
| Dyslipidemia | 0 (0%) | 5 (45%) | 0.12 |
| Hypertension | 2 (40%) | 8 (72%) | 0.30 |

Fisher's exact test or Mann-Whiteney U test
