## Supplemental Table 2 for "Neutrophil extracellular traps and thrombotic components across human coronary plaque progression: Association of intraplaque hemorrhage with NET formation"

**Supplementary table 2. Immunopositive area for cellular and molecular components in human coronary artery**

| | Diffuse intinal thickeing (n=23) | Pathological intinal thickening (n=26) | fibrous cap atheroma (n=14) | Fibrocalcified plaque (n=16) | Thin-cap fibroatheroma (n=10) | Ruptured plaque (n=13) |
|---|---|---|---|---|---|---|
| SMA (%) | 13.7 (11.4-19.3) | 6.6 (4.05-12.18) | 3.8 (1.05-5.4) | 3.75 (2.33-5.28) | 0.2 (0-0.55) | 0.46 (0.03-0.86) |
| CD68 (%) | 0.01 (0-0.03) | 0.045 (0.01-0.17) | 1.42 (1.06-2.99) | 0.21 (0.06-0.57) | 4.65 (2.4-6.15) | 3.7 (2.32-7.84) |
| CD66b (%) | 0 (0-0) | 0 (0-0) | 0 (0-0.023) | 0 (0-0) | 0.03 (0-0.2) | 2.1 (0.32-6.8) |
| citrullinated histone H3 (%) | 0 (0-0) | 0 (0-0) | 0 (0-0.04) | 0 (0-0) | 0.05 (0.03-0.12) | 0.17 (0.01-1.07) |
| glycoprotein IIb/IIIa (%) | 0 (0-0) | 0 (0-0) | 0 (0-0) | 0 (0-0) | 0.01 (0-0.05) | 0.37 (0.12-1.31) |
| fibrin (%) | 0 (0-0) | 0 (0-0.08) | 0.79 (0-3.83) | 0 (0-0.22) | 6.36 (5.12-11.82) | 14.21 (5.71-21) |
| von Willebrand factor (%) | 0 (0-0) | 0 (0-0) | 0 (0-0.14) | 0 (0-0.03) | 0.41 (0.06-2.62) | 11 (7.3-22) |

Date are expressed as median and interquative range
